# Auditory feedback effect of self-pacing treadmill walking in comparison to overground walking

**DOI:** 10.1101/2024.08.28.610096

**Authors:** Trevor V. Evans, Megan E. Reissman, Timothy Reissman, Mark Shelhamer, Ajit M. W. Chaudhari

## Abstract

Self-pacing treadmills provide advantages for assessing locomotion, but there are differences between gait patterns in overground and treadmill paradigms. A potential explanation could be that the variable auditory sound from the treadmill belt motors when changing speed provides artificial sensory feedback to walkers that may influence their motor output while on the self-pacing treadmill. We hypothesized that there would be significantly different temporal measures between the sound and no-sound conditions, and these same measures from the no-sound condition would be similar to overground walking. Participants (n=29) walked under three different conditions for five-minute periods each: on the self-pacing treadmill with sound inclusion, on the self-pacing treadmill with sound exclusion, and overground walking. Significant differences were found in the variability of all temporal walking measurements between the overground and both treadmill conditions (p < 0.05). Effects of sound on temporal walking measurements were not observed. The lack of difference between the two treadmill conditions suggests that self-pacing treadmill walkers may not utilize the variable belt motor sounds available to them. Additionally, overground walking still has different gait patterns compared to self-pacing treadmill walking, so other potential sources of differences should be explored in future work. Given that both treadmill conditions had different temporal gait patterns from overground, considerations for using the self-pacing treadmill as an alternative for overground walking should be taken when studying human locomotion.

## Introduction

### Overground vs treadmill walking

Treadmills are frequently utilized in research for gait analysis to simulate overground locomotion ^1–3^. They offer several advantages including: controlled indoor environments, convenience to walk in a relatively small space, and ability to collect very accurate biomechanical data over long time periods. Standard treadmills operate at one speed that is pre-determined before the belts begin to move or require manual input to change the belt speed. Walking on a fixed-speed treadmill (FST) does not allow for spontaneous speed fluctuations, which we know occur in standard overground walking ^4,5^. Fixing a key aspect, such as gait speed, also likely affects spatiotemporal parameters such as stride time and stride length which are directly related to speed ^6^. This limits the generalization of FST walking research to human locomotion. Other studies comparing overground to FST walking have found differences in kinematic, spatiotemporal, and kinetic parameters ^7^.

The development of a self-pacing treadmill (SPT) enables continuously adjusting belt speed so that a preferred speed no longer must be predetermined and fixed, while retaining the FST advantages ^8,9^. Several feedback control approaches for SPT have been identified, including position-based ^8,10–14^, behavior-based ^15^, and force-based ^16–20^ algorithms. Only a few studies have compared SPT and overground walking paradigms in healthy adult populations ^10,21,11^, which is important for validation in future research for drawing conclusions from gait analysis done on SPT. Gait speed, often regarded as the most important measure of gait, was not found to have significant differences between SPT and overground walking ^10,21^. Spatiotemporal measures show mixed findings since one study reported differences in swing and stance, single and double support, and step width among others ^21^, while another study reported no difference among any of the variables ^11^. Both study protocols used position-based feedback control and collected overground data in similar ways, so there remains a need to further research why spatiotemporal differences may exist between the two walking paradigms.

### Role of sound in movement

SPT belts are actuated by large motors, which emit auditory noise at variable intensity and frequency levels dependent on the belt’s instantaneous velocity and acceleration. The exact auditory characteristics change based on the hardware design of the treadmill. These sounds are perceptible stimuli for the person walking on the SPT, who is constantly using several sensory feedback modalities to maintain their balance during walking ^22,23^.

Hearing allows us to perceive sound information from our environment, and we incorporate auditory stimuli among our other senses to aid in making goal-oriented decisions ^24,25^. In walking, we use hearing to receive spatial and temporal cues that may help better control our balance ^26^. External perceivable auditory cues have been shown to alter gait performance in both healthy ^27,28^ and Parkinson’s disease ^29,30^ populations. One study reduced auditory feedback during gait using ear plugs and white noise in older adults, but found no differences in spatial or temporal aspects of walking ^31^. However, they used a FST which does not have the variable sound profile that a SPT does. When this feedback is removed, temporal parameters from this SPT paradigm could be closer to measurements observed during overground walking.

### Purpose

The purpose of this study was to evaluate the effect of audible treadmill motor belt sounds on self-pacing treadmill walking patterns. We hypothesized that significant differences in temporal gait characteristics would exist between conditions where participants could hear the variable sounds from the self-pacing treadmill belt motors versus when they could not. Furthermore, we believed that these same characteristics would be similar between the no-sound and overground conditions.

## Methods

### Participant recruitment

All study procedures were approved by The Ohio State University Institutional Review Board prior to enrolling any participants. Healthy adult participants within the age range 18-55 in the local area surrounding The Ohio State University were recruited for this study. Recruitment began on the 29^th^ of March, 2022 and lasted until the 19^th^ of February, 2024. Paper flyers and email listservs served as the means for recruitment. Participants who were unable to walk for an hour consecutively, or who had a diagnosed lower body injury within the past three months from time of collection, were excluded from the study. Those who met the criteria were enrolled in the study as participants after providing institutional review board–approved written informed consent.

### Experimental protocol

The protocol outlined and data collected here were a part of a larger study. The following are details for one of two portions of the study in which this details the overground portion. Wireless tri-axial accelerometer sensors (Trigno Legacy, Delsys Inc.; Natick, MA) were placed at three locations along each participant’s body: one directly below each of the left and right medial tibial condyles on the shank, and one over the sacrum. The leg sensors were secured using Velcro wraps, and the sacral sensor was placed in a pouch inside of a belt around the lower back. Participants walked on an indoor 80-meter track above the lab and completed one six-minute recorded trial at their preferred pace. The first lap was not recorded to ensure the trials only captured gait cycles around steady-state speed and to maintain starting point consistency across participants.

The following are details for the second of two portions of the study in which this details the self-pacing treadmill (SPT) portion. The accelerometer sensors remained on for this portion. A lower back cluster of reflective markers was placed over the sacrum. Functional joint center and bony landmark calibrations were performed to properly scale the internal skeletal model based on the marker cluster placement. The marker cluster was used to determine the walker’s fore-aft position along the treadmill. Participants were attached to an overhead safety harness on an instrumented split-belt treadmill (Immersive Lab; Bertec Corp.; Columbus, OH), given a brief explanation of how a SPT functions differently from a fixed-speed treadmill, and instructed to walk at their preferred speed for the duration of the protocol. The treadmill used a positional-based control to modulate belt speed. Participants began to walk for an acclimation period of at least five minutes, after which they were asked to confirm that they felt comfortable with walking on the SPT. The baseline SPT walking trial consisted of one five-minute recording period of accelerometer data while hearing all environmental sounds.

Participants were then given noise-cancelling wireless headphones (Model QuietComfort 35; Bose Corp., Framingham, MA) to remove perceived audible treadmill belt motor sound from the participant while walking. Brown noise ^32^ was played through the headphones, with loudness level increasing incrementally until participants confirmed they could not hear any environmental sound. Accelerometer data from one five-minute SPT walking trial with headphones worn were collected.

The self-pacing treadmill controller used for both treadmill walking trials was created and executed in MotionMonitor xGen (Version 3.57; Innovative Sports Training; Chicago, IL). MotionMonitor’s built-in proportional feedback controller was used to adjust the belt speed based on the anterior-posterior position of the lower back marker cluster relative to the center of the treadmill. Marker trajectories were tracked in real-time using MotionMonitor connected to a nine-camera Vicon motion capture setup (Tracker Software, Vero Cameras; Vicon Motion Systems Ltd; Oxford, UK).

Three-axis accelerometer data from all three walking trials (treadmill baseline, treadmill with sound removal, overground) of the left shank, right shank, and pelvis sensors were captured at 148 Hz and saved in MotionMonitor. Vertical ground reaction force data recorded at 1000 Hz from the treadmill walking trials was also captured and saved in MotionMonitor.

Accelerometer data were analyzed in MATLAB using custom written scripts (Version 2023b; MathWorks, Inc.; Natick, MA). Ground reaction force data were analyzed in MATLAB using custom written scripts.

### Gait event detection

Gait event detection is important for identifying events used to characterize human walking, particularly heel-strike and toe-off times. Wearable inertial sensors have recently been favored due to their compact and inexpensive designs, and event detection using them shows low error in comparison to the standard force plate and foot switch equipment (Lee et al, 2007; Mansour et al, 2015; Gurchiek et al, 2020). Therefore, these sensors would be appropriate to assess gait events during overground walking, where there is a lack of data in the literature.

The algorithm used to identify the events is based on a previous study that detected gait events from thigh-worn accelerometers ^33^. Gait events here were calculated from accelerometer data of the sensors placed on the medial left and right shanks. Left heel-strike and toe-off events were found using data from the long axis of the left shank, which is oriented vertically when participants stood in the anatomical position. First, single-sided power spectral densities (PSD) of the accelerometer signal were estimated using Fast Fourier Transformations. A moving average filter with window size of 50 was applied to the PSD data to reduce noise and identify the two largest frequencies in the signal. The frequency at which the largest peak between 0.6 – 1.4 Hz occurred was estimated as the stride frequency (f_str, L_). The largest peak in the range of 1.4 – 2.5 Hz was estimated as the step frequency (f_stp, L_). The accelerometer signal was then low-pass filtered at three cutoff frequencies: f_str_, f_stp_, and 5 x f_str_, with the last of the frequencies being chosen to remove noise while maintaining fidelity of the true movement data. Using the stride frequency to estimate the largest cutoff frequency allows for participant-specific adaptability to a range of gait speeds. In total, three filtered accelerometer signals were used to estimate gait events for the left shank (Fig 1).

**Fig 1.**
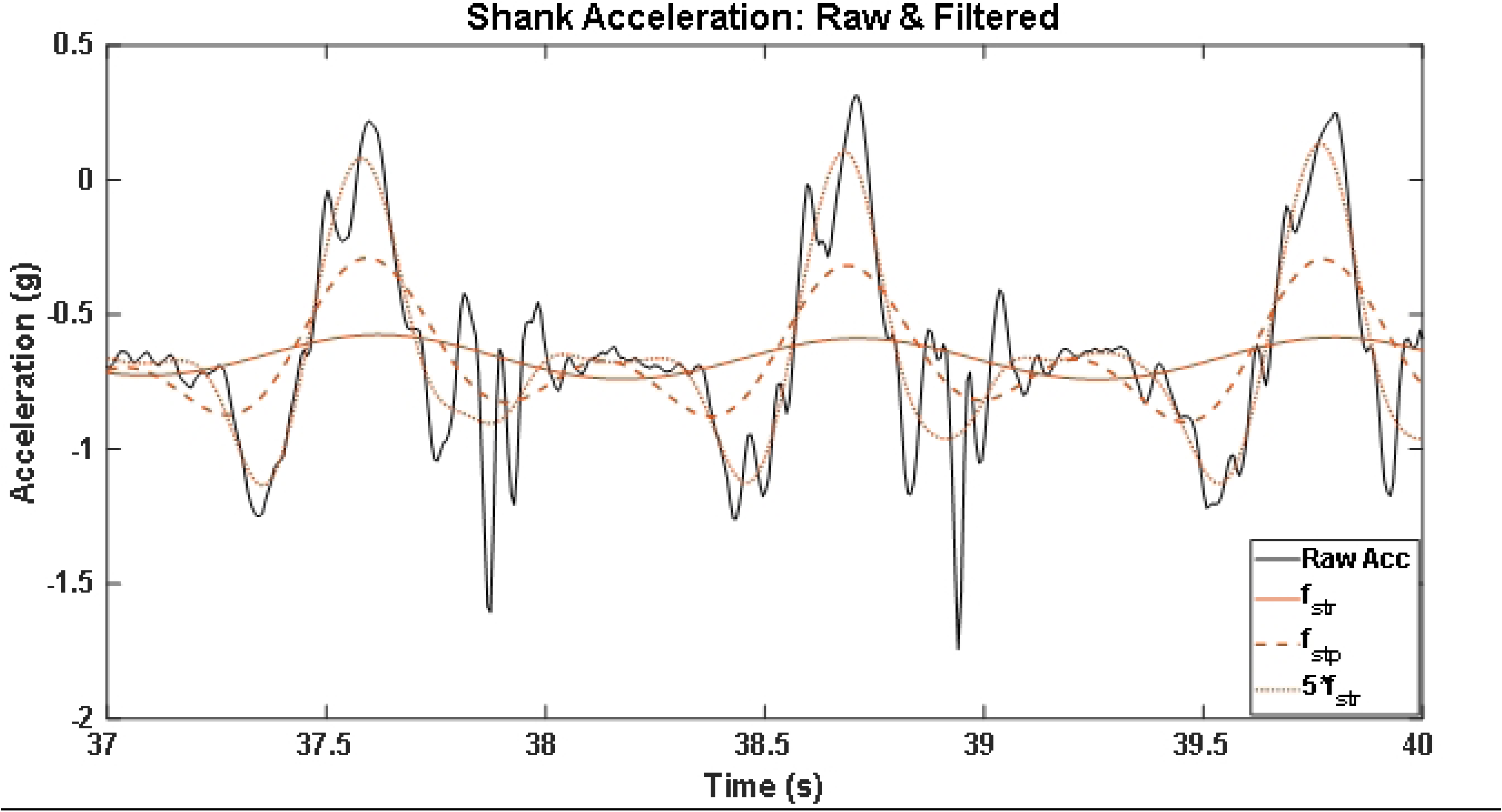
Shank acceleration: raw & filtered. Raw long-axis shank acceleration (black solid) and filtered shank acceleration at three frequencies: stride frequency (f_str_; solid orange), step frequency (f_stp_; thick dashed orange), and 5 x stride frequency (5 x f_str_; thin dashed orange) of baseline.

To identify gait events from accelerometer signal features, all maxima of the f_str_-filtered accelerometer signal were first identified (Fig 2, Arrow 1), which were general indicators of the contralateral leg being in the mid-swing phase. Next, the minimum value of the 5 x f_str_ signal directly preceding each f_str_ maximum was estimated as the toe-off event (green triangle) (Fig 2, Arrow 2). For heel-strike identification, the succeeding crossing of the f_str_-filtered signal with the f_stp_-filtered signal after each f_str_ maximum was identified and chosen as the estimate (Fig 2, Arrow 3). This process was repeated for the right shank accelerometer data to detect gait events.

**Fig 2.**
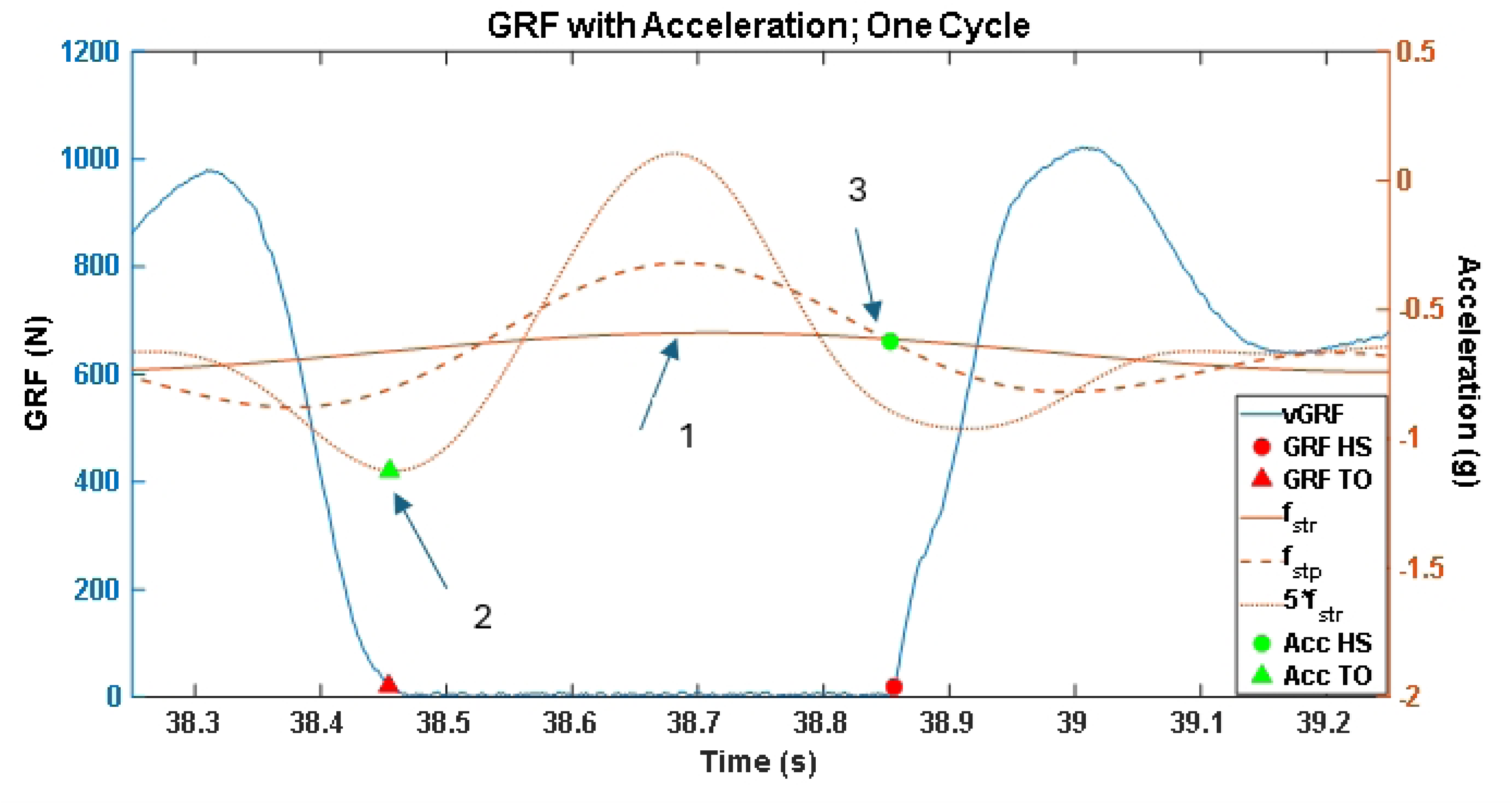
vGRF and filtered acceleration signals with detected gait events. ; Arrow 1: maximum values of the f_str_ signal; Arrow 2 – The time at which the minimum value of the 5 x f_str_ signal directly preceding each f_str_ maximum is estimated as the toe-off event (green triangle); Arrow 3 – Following each f_str_ maximum, the intersection of the f_str_ and f_stp_ curve is estimated as the heel-strike event (green circle)

To validate this event-detection algorithm, vertical ground reaction force (GRF) data from the baseline treadmill walking trial were used as baseline reference values. A 20 N threshold was used to identify the heel-strike and toe-off events (Fig 3). Positive crossings were identified as heel-strike events, and negative crossings were identified as toe-off events.

**Fig 3.**
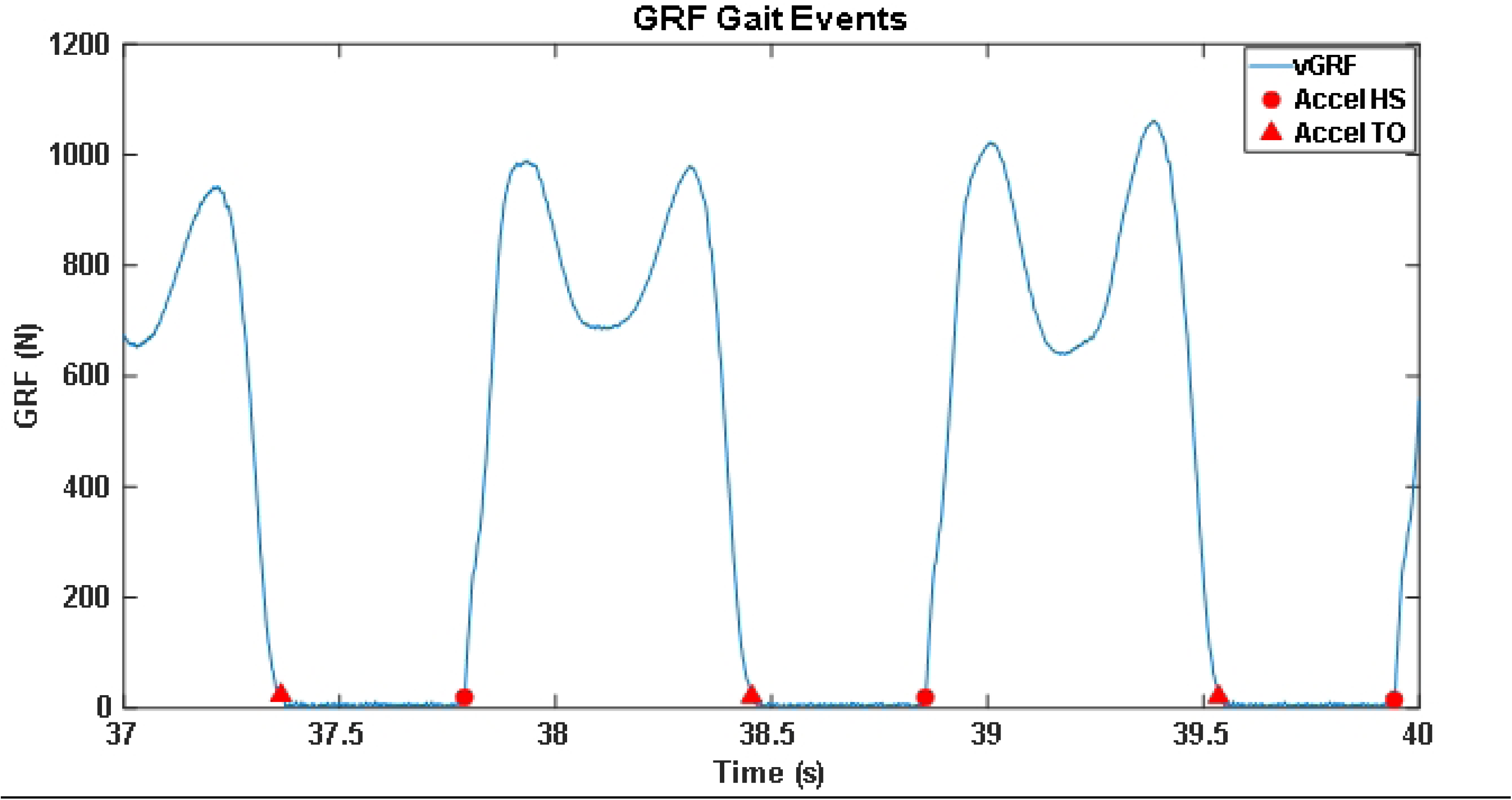
Event detection from vGRF validation data.

Twelve participants were used for the validation of the accelerometer event detection algorithm out of the total 34 who enrolled in the study. Validation was performed based on a the five-minute duration baseline walking condition. Seventeen participants were excluded from the validation analysis because they placed both feet on the same belt for a significant portion of at least one of the treadmill walking trials, resulting in no usable GRF data. Another five participants were excluded from both the algorithm validation and the condition comparison analysis due to technical difficulties.

### Statistical analysis

Accuracy of the gait event detection algorithm was quantified in MATLAB by calculating the absolute error of the accelerometer-based timing estimates from the reference GRF-based times of heel-strike and toe-off events.

Linear mixed models for repeated measures with walking paradigm as the fixed effect and participants as random effects were used to find changes in both the average and standard deviation of various temporal parameters (JMP Pro 17.0; JMP Statistical Discovery LLC; Cary, NC). The significance level was set at α = 0.05. Post-hoc Tukey Honest Significant Differences tests were conducted to make pairwise comparisons when appropriate.

## Results

### Accelerometer event detection algorithm validation

Participants (n=34) were enrolled in the study that satisfied the inclusion criteria. Data from 12 of these participants (4 M, 8 F; Age: 22.7 ± 4.16 yrs.) were used to validate the accelerometer event detection algorithm by comparing it to reference ground reaction force data.

Table 1 summarizes the errors in the accelerometer-detected gait events compared to reference values from the ground reaction force data. An average of 535 steps from each participant during their five-minute baseline SPT walking trial was used to validate the algorithm. The average absolute errors for heel-strike events across all participants were 15.9 ms and 20.8 ms for the left and right foot, respectively. The average absolute errors for toe-off events across all participants were 22.4 ms and 19.3 ms for the left and right foot, respectively.

**Table 1.** Event detection error differences. Error between gait events detected from shank accelerometers and force plates; SD: standard deviation averaged across all participants.

|  |  | Gait Event |  |  |  |
| --- | --- | --- | --- | --- | --- |
|  |  | Heel Strike |  | Toe Off |  |
|  |  | Left | Right | Left | Right |
| Error | Mean Abs. Error ± SD (ms) | 15.9 ± 11.6 | 20.8 ± 16.5 | 22.4 ± 14.4 | 19.3 ± 12.6 |

Table 1
|  |  | Gait Event |  |  |  |
| --- | --- | --- | --- | --- | --- |
|  |  | Heel Strike |  | Toe Off |  |
|  |  | Left | Right | Left | Right |
| Error | Mean Abs. Error ±<br>SD (<u>ms</u>) | 15.9 ± 11.6 | 20.8 ± 16.5 | 22.4 ± 14.4 | 19.3 ± 12.6 |

### Walking paradigm comparison

Data from 29 participants (15 M, 14 F; Age: 21.9± 3.14 yrs.) were used to compare the three walking paradigms (SPT with sound, SPT with sound removed, and overground) after eliminating data from five participants (n=4 stopped walking during one of the three trials, n=1 had sensor issues).

The linear mixed effects model showed significant main effect differences in variability between walking conditions (stride time: p < 0.0001, step time: p = 0.0007, stance time: p < 0.0001, swing time: p = 0.0002). Post-hoc pairwise comparisons showed significant increases in variability during the overground condition compared to the SPT without sound conditions (stride time: p < 0.0001, step time: p = 0.0007, stance time: p < 0.0001, swing time: p = 0.0003) and overground compared to SPT with sound conditions (stride time: p = 0.002, step time: p = 0.0107, stance time: p < 0.0001, swing time: p = 0.002). No significant differences were found between the SPT sound compared to the no sound conditions (stride time: p = 0.3495, step time: p = 0.6491, stance time: p = 0.94, swing time: p = 0.82), indicating that external auditory sound had no statistical significance on the variability of temporal measurements collected during SPT walking. Fig 4-7 summarizes the comparison of temporal variability (standard deviation) measurements of stride time, step time, swing time, and stance time between the three walking conditions.

**Fig 4.**
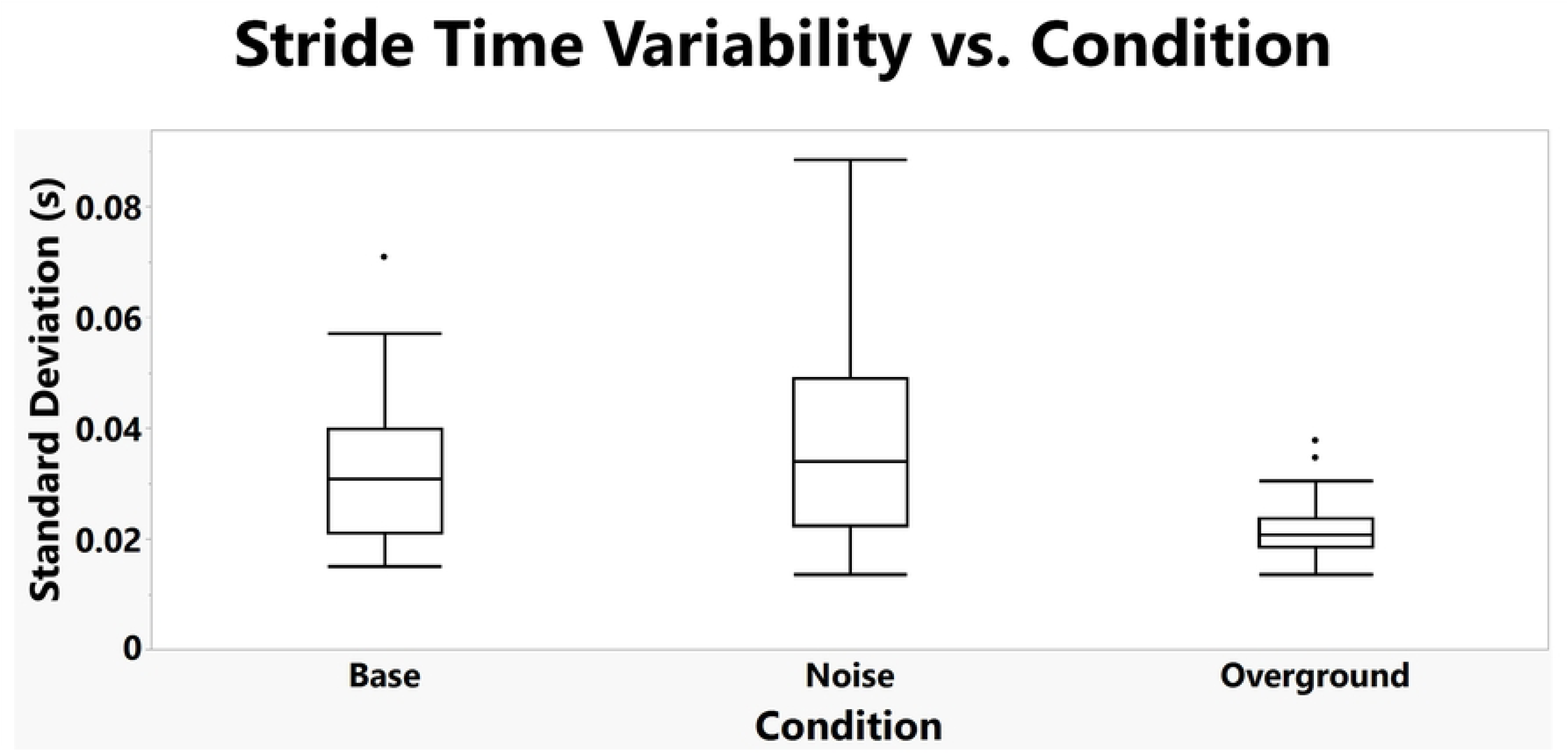
Stride Time Variability across walking conditions.

**Fig 5.**
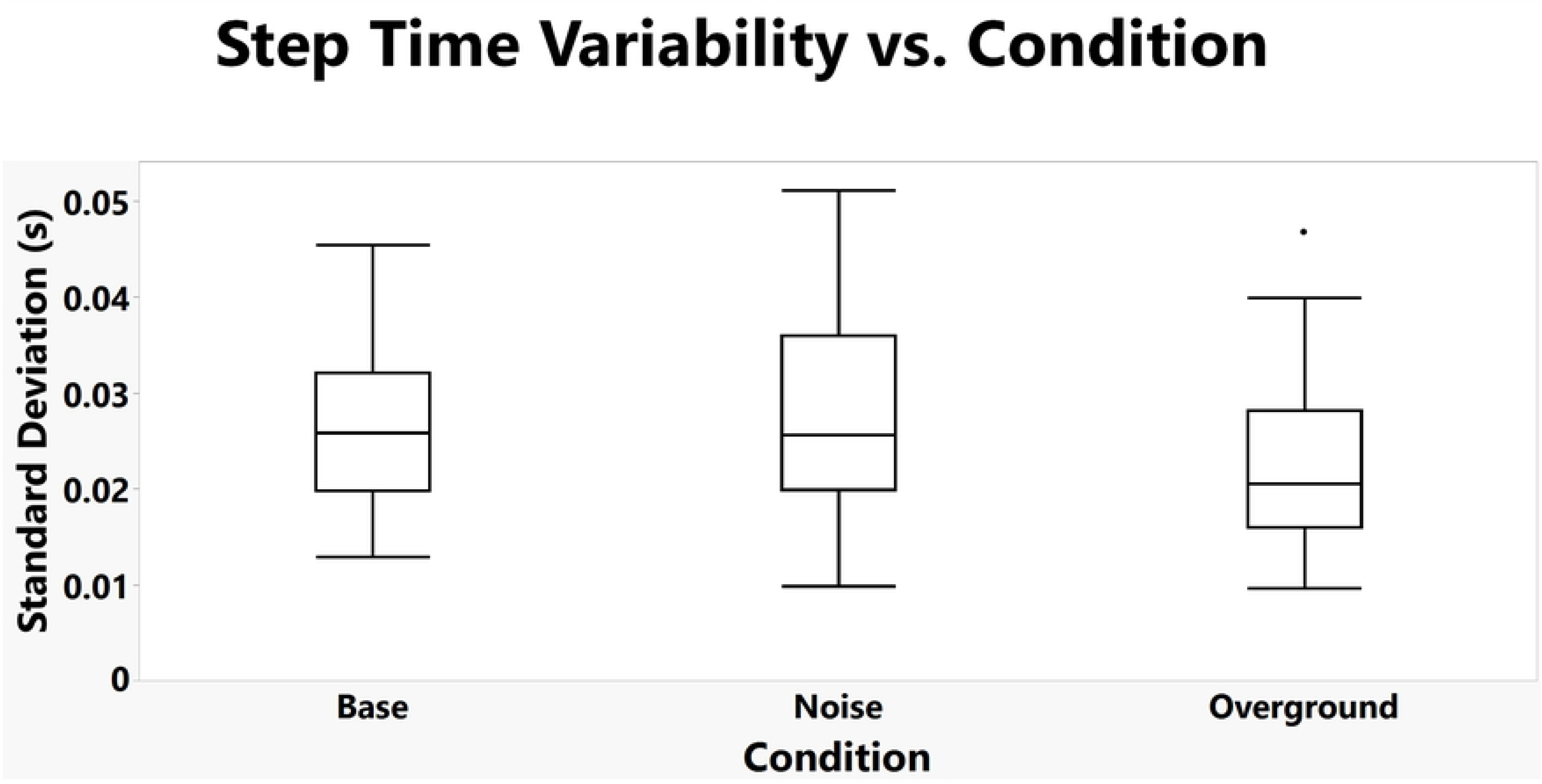
Step Time Variability across walking conditions.

**Fig 6.**
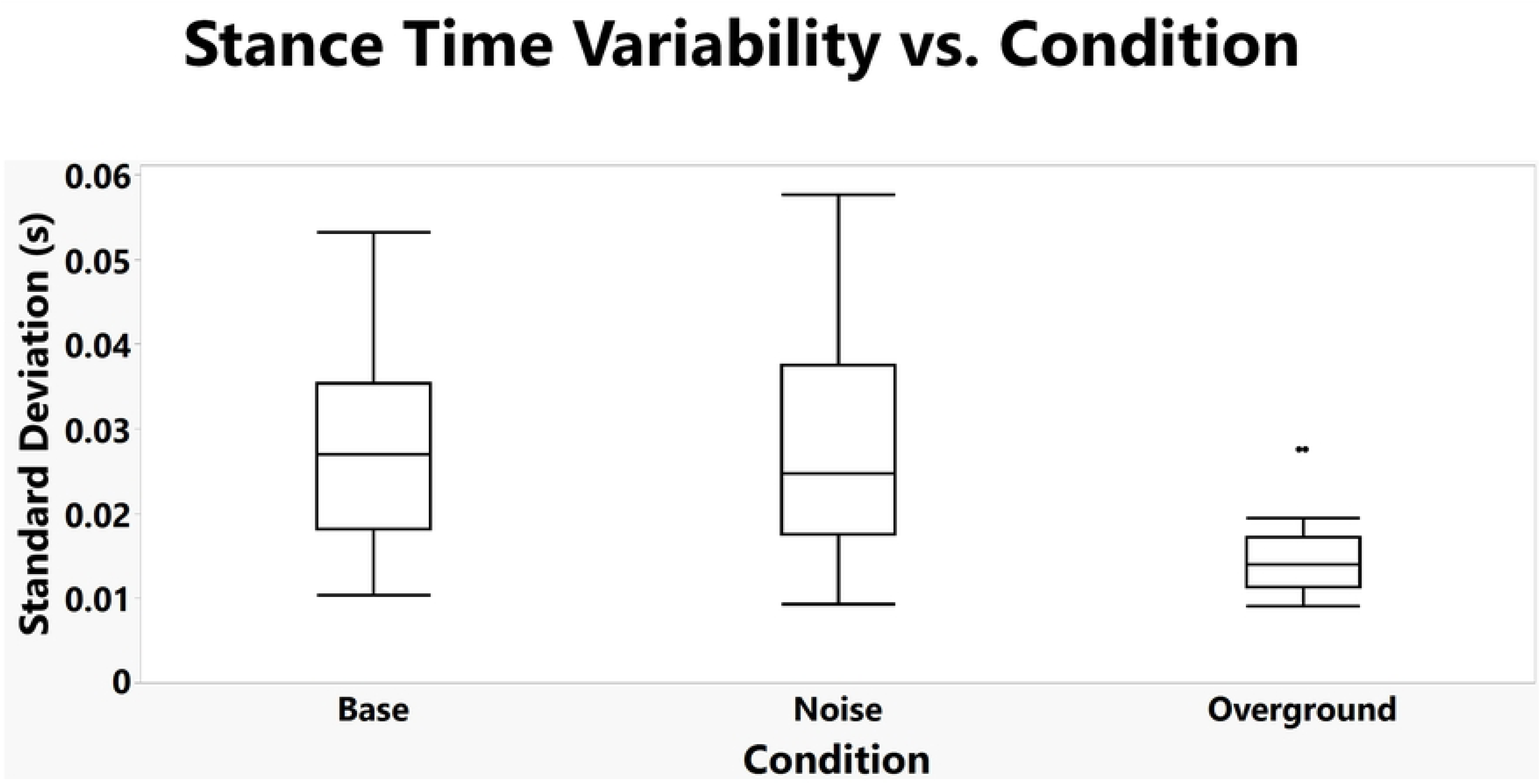
Stance Time Variability across walking conditions.

**Fig 7.**
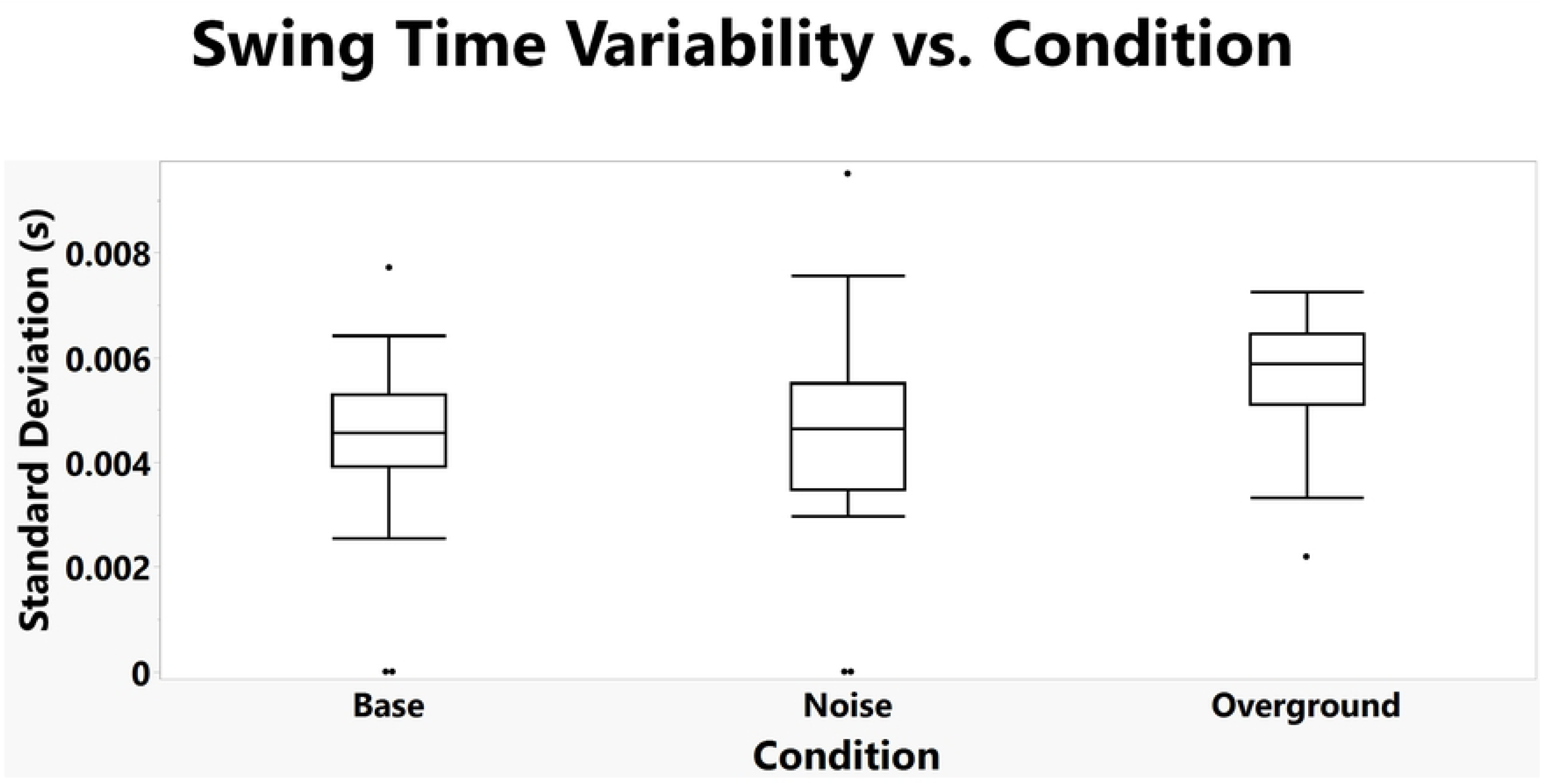
Swing Time Variability across walking conditions.

Significant differences in mean values between conditions were not found for any of the temporal measures (stride time: p = 0.184, step time: p = 0.1863, stance time: p = 0.1474, swing time: p = 0.358), indicating that there were no statistically significant differences in the mean values of temporal measurements collected during all three walking conditions.

Fig 8 shows each participant’s average stride time for each of the three walking conditions. No trends in individual average stride times were identified across conditions.

**Fig 8.**
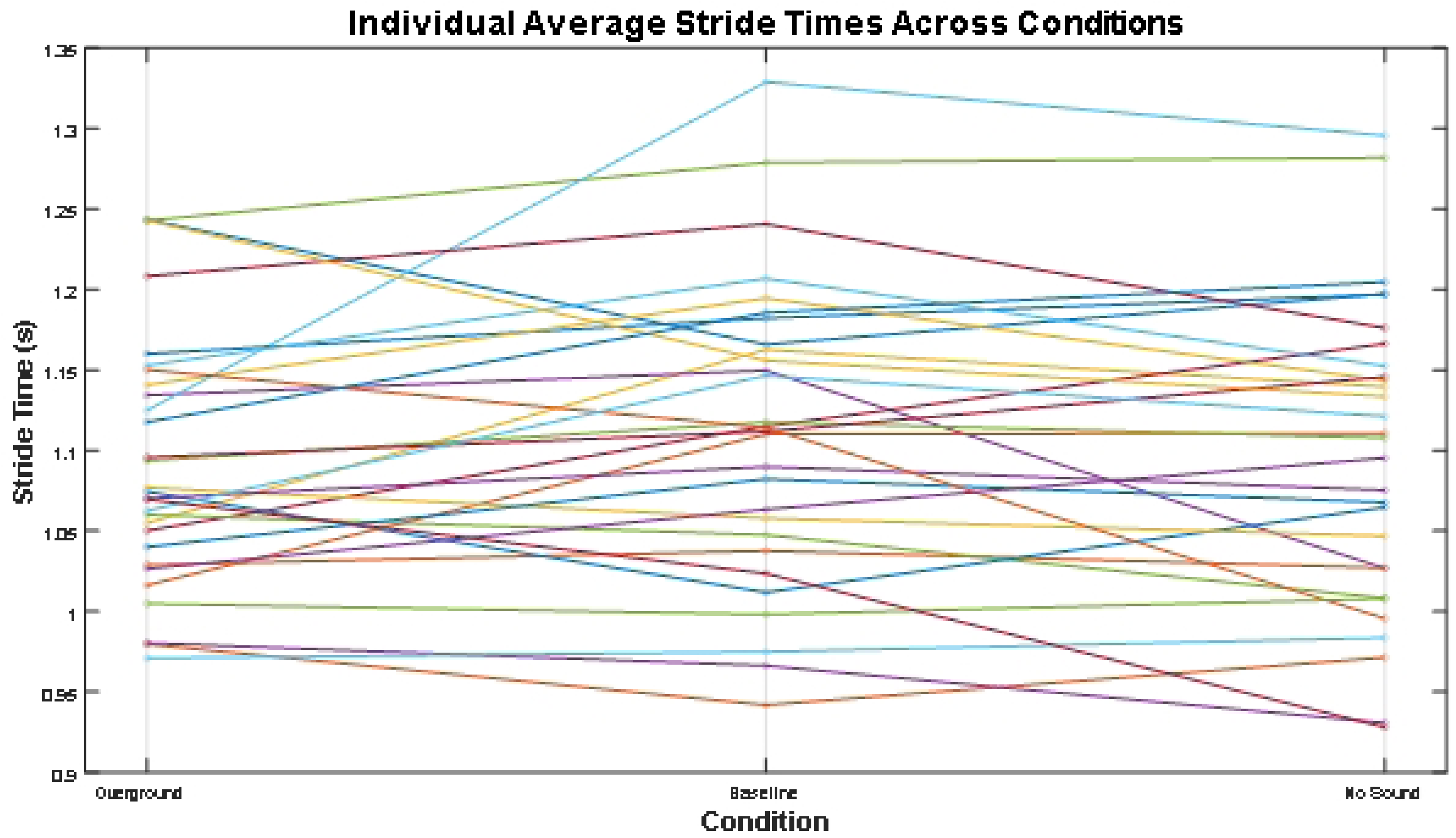
Individual Average Stride Time across walking conditions.

## Discussion

This study explored the effect of auditory feedback during self-pacing treadmill (SPT) walking on gait parameters compared to overground walking. We hypothesized that there would be significant differences in temporal walking patterns from baseline when auditory feedback from the treadmill belt motors was removed. Furthermore, we hypothesized that removing the feedback would lead to gait patterns that more closely resembled overground walking. Neither of these hypotheses was supported, as we saw no differences between the two treadmill conditions. No differences were found in temporal measurements between the baseline condition with treadmill sound and the brown noise condition that removed external sound. Significant differences were seen when comparing both treadmill walking conditions to overground walking. These results are consistent with some previous work, which found significant differences between SPT and overground walking paradigms in stride time, swing and stance percentage, double and single support percentage, among other temporal measurements ^21^. In contrast, Reneaud et al. ^11^ found a lack of differences in stride time, stance phase, and cadence measures. This discrepancy could be for several reasons, one being that they collected their overground data from walking back and forth in a small corridor, so only twelve gait cycles were used for analysis. Another is that on a small corridor, steady-state walking may not be reached before the person must stop to switch their walking direction. Additionally, it has been observed that the type of treadmill controller used for SPT operation affects gait patterns ^34,35^, so the aforementioned studies with conflicting results may have utilized different controllers. This study used positional control where the belt speed changed proportional to the fore-aft distance of a sacral marker from the center of the treadmill.

These results suggest that walkers on a SPT do not process the belt motor sounds as useful feedback to plan and initiate their movements on the treadmill. One reason that participants did not respond to the sound feedback may have been delays in sound feedback relative to other cues such as visual ones. Recall that when the participant walks and their estimated center of mass crosses the treadmill’s fore-aft midpoint, the controller triggers the motor to accelerate or decelerate the belts proportional to the distance of the walker’s center of mass from the treadmill’s midpoint. Given the treadmill’s inertia, there is a delay between the trigger point and the treadmill reaching its desired speed based on COM displacement. During that delay, however, the belt motors provide the auditory feedback to inform the walker of the forthcoming adjustment of the treadmill belts. The feedback serves two purposes: it informs the walker about the imminent adjustment and provides spatial information about their relative position, which could be important in a treadmill environment due to a lack of visual cues which are known to be important for walking ^34,36^. However, SPT walkers might adapt to a lack of optic flow and other auditory cues since movement and spatial processing are multisensory responsibilities ^37–39^. This may emphasize the need for familiarization periods when walking on a self-pacing treadmill ^40^. Incorporation of the sound from the motors may be relevant for pathological or older populations who have diminished sensory capabilities or a greater fear of falling. An additional sensory modality during walking could aid in kinesthesia and environmental awareness, alleviating the poorer quality of information that comes with age and disease. More research needs to be done to fully understand the various sensory contributions and reweighting performed while walking on a SPT.

While there were no significant differences in the temporal measures between treadmill conditions, the treadmill conditions exhibited significantly higher variability in stride, step, and stance time compared to the overground condition. This could be a function of walking at a lower speed than typically preferred, which has been associated with higher gait variability ^41–44^. Decreased speed and increased variability are typical aspects of a cautious gait pattern, which participants may have demonstrated given the novelty of walking on a self-pacing treadmill. However, gait speed data were not collected for this study since the sensors used for data collection were unable to accurately estimate walking speed. Past work has shown gait variability is influenced by environment and walking conditions ^45^. This suggests that the variability increases may be attributed to the new experience and potential discomfort of walking on a self-pacing treadmill, regardless of the inclusion or exclusion of auditory feedback from the treadmill belt motors. This idea warrants further research into how motor control strategies may differ between treadmill and overground walking.

While significant differences were found in variability measures, there were no differences found in mean temporal measures between conditions. This indicates that walking condition had no effect on the mean values of the calculated measures, even if those measures may have had large fluctuations throughout the trials.

This study also presents a novel algorithm for detecting gait events from shank accelerometers on both legs. The algorithm developed here is inspired by previous work ^33^ but improves upon the errors calculated from their thigh-worn accelerometers. The absolute errors found presently are comparable or less than the values stated there, suggesting that this method may be a more accurate estimator of gait events when walking in both treadmill and overground environments. We found errors of < 23 ms, which was about 2% of the gait cycle on average. This contradicts the findings in ^46^ which it was claimed that shank accelerometers were inferior placements for wearable IMUs to detect gait events, since they found errors of up to 80 ms when comparing event times to gold standard force plate data. Additionally, this study provided a thorough dataset lacking in the literature to characterize overground walking, particularly in spaces where participants can take large numbers of continuous strides for gait assessment.

However, a few limitations exist. The ordering of the trials was not randomized; the sequence was overground ◊ treadmill baseline ◊ treadmill without auditory feedback for all participants. This order was chosen because a full-body marker set and other equipment were placed on participants for the treadmill trials as this study was part of a larger project, so it made sense to do the overground condition first. The fixed ordering could introduce various types of biases to the study that confound the results, and any conclusions drawn may not actually answer the desired research questions. This may include learning effects, fatigue, or selection bias. However, since overground walking is how humans naturally walk, using that condition first should not introduce any learning effects. To our knowledge, no studies have found any carry-over effects of overground walking before treadmill walking. Additionally, sufficient time was waited between conditions to avoid order bias issues. Another limitation is that the two treadmill conditions compared here were not always done consecutively. Since this was part of a larger walking study, there were sometimes other activities participants between the baseline and headphone conditions analyzed here. This intervention could possibly confound the results. This is unlikely though, because at least five minutes of wash-out between every condition for the study was done to minimize any bias. Since people are likely more accustomed to walking on a treadmill while able to hear external sounds, putting the sound exclusion trial last will not be an issue since that condition is the most novel. Additionally, the accelerometers used for the study were slightly off-axis of the typical expected coordinate system of an IMU. For example, standing the sensor straight up did not correspond to exactly ±1 g on the long axis of the sensor, which could impact the transfer of the developed algorithm to other properly calibrated sensors. However, this is unlikely since our pilot testing showed that the sensors used still reliably possess the same general kinematic patterns during gait. Further research is needed to validate the event detection algorithm in clinical populations and in daily movements other than steady-state walking. Walking typically involves optic flow, however it is constrained by the length of the belts on a fixed reference frame treadmill. Additionally, walkers faced a gray screen while walking, which further reduced any potential optic flow, which may have impacted their ability to assess their orientation and movement in space. Optic flow affects gait variability ^36,47–49^, so that could explain the differences seen between treadmill and overground walking.

Only temporal gait measures were obtained here due to equipment restrictions for the overground walking condition. Thus, we were unable to assess whether spatial, kinematic, and kinetic parameters were affected by the auditory feedback from the belt motors on the treadmill, and if that would have resulted in similar overground walking values.

Brown noise played through worn headphones during the sound exclusion condition was loud enough to block the variable treadmill motor belt sounds while walking. This was effective as it blocked all environmental sounds from being heard. This includes the sound of footsteps that impact walking parameters ^50,31^, which may be heightened on a treadmill due to the louder sound from frictional forces during heel strike. The exclusion of the treadmill belt motors could have also blocked natural footstep sounds, which may be a confounding factor when interpreting significant results between the treadmill and overground conditions.

Finally, considerations for the exact treadmill characteristics should be taken. The instrumented treadmill used in this study was approximately 1.8 m long, so future work should explore potential effects of SPT belt length on gait patterns. Additionally, other SPT may have different sound profiles that could affect auditory perception and ultimately, gait patterns.

## Conclusions

Variable treadmill belt motor sounds were found not to significantly affect temporal measures during walking. Participants walking overground, on the treadmill without headphones, and on the treadmill with headphones demonstrated similar gait patterns, suggesting that these sounds are not used as auditory feedback for motor planning and control on the treadmill. Additionally, an accelerometer-based gait event detection algorithm was developed and validated with high accuracy. The algorithm can be used for future work to study gait, and further investigation into self-pacing treadmills is necessary to fully understand their differences from overground walking conditions.

## Acknowledgements

We are thankful to Dan Merfeld for his helpful comments with this study, as well as Steve Wilson and Raymon Patton for their assistance during data collection. We are also appreciative of the study participants for volunteering. Trevor Evans was supported for this study from MURI #12731926.

